# Insights into structures of peptide aggregation nuclei from concentration dependence of lag time

**DOI:** 10.1101/2025.09.19.677452

**Authors:** Egor O. Vasilenko, Dmitry N. Ivankov

## Abstract

Many peptides and proteins self-assemble into large fibrillar aggregates, reaching sizes of several micrometers. This process typically involves nucleation, formation of transient oligomeric species ranging from dimers to assemblies comprising hundreds of monomers. The roles of these heterogeneous oligomers in initiating fibril growth vary significantly, as only a subset converts into primary nuclei, the smallest assemblies capable of spontaneous elongation. Consequently, the initial stages of peptide aggregation remain a critical challenge in elucidating amyloid fibril formation. Here, we analyze the aggregation lag time as a function of initial peptide concentration, employing linear regression on a double logarithmic scale to derive the critical nucleus size from the slope. We compare potential shifts in kinetic regimes with critical concentrations associated with the assembly of larger oligomers. Although peptide length has no clear correlation with nucleus size, extended helical regions may promote the formation of larger nuclei, stabilizing preliminary oligomers and prolonging the lag phase. We quantified the contributions of higher-order oligomers as on- and off-pathway species across multiple peptides, identifying a common critical micelle concentration range. We prove that linear growth models can not capture the weak concentration dependence of lag times for several peptides. We demonstrate that the inclusion of capping and fragmentation mechanisms substantially improves the plausibility of the model. Knowledge of nucleus size facilitates molecular dynamics simulations to capture transitions to fibril-prone conformations. The insights advance our understanding of amyloid nucleation, identifying toxic aggregates in human neuroglial cells, and research on drug safety and biotechnological applications.

## 1 Introduction

Alzheimer’s [1] and other neurodegenerative diseases [2], cataract formation [3], and many other serious disorders are associated with peptide aggregation. This phenomenon is dramatically accelerated in the presence of seeds (nuclei) [3, 4]. This means that the primary assembly of nuclei is of critical importance for the subsequent growth into large, prolonged fibrils or amorphous granules [1, 3, 5]. Therefore, the development of quantitative models of nucleation has significant potential to enhance preventive and therapeutic pharmaceutical strategies [1, 4], improve safety validation of prospective drugs [4], and facilitate the development of therapeutic interventions and preventive measures against pathological aggregation. At the same time, seed-based strategies are emerging as a promising therapeutic approach against neurodegenerative diseases and diabetes [4]. Peptide and protein aggregation can sometimes be functional [1] as in the case of microfilaments [6], storage hubs, and catalytic surfaces within the cell [2]. Algorithms that predict growth rates from a peptide’s molecular properties could significantly reduce resource expenditure in nanotechnology industries [6] and improve the efficiency of protein crystallization for the determination of tertiary structures [7]. The presence of diverse absorbance, scattering, and fluorescence kinetic time profiles makes it possible to validate theoretical models [1, 3, 5, 8, 9]. Comparing various peptides to obtain evidence on general relationships between primary and secondary peptide structures and nucleation propensity may facilitate drug developments and allow extrapolation of prediction algorithms to peptide-lipid and peptide-nucleic acid interactions, which are often the key events for their physiological roles.

Functions representing aggregation rates are typically constructed from linear and exponential terms, accounting for the contributions of primary and secondary nucleation, branching, and fragmentation [10]. Many general growth functions, such as the widely employed Finke-Watzky mechanism [3, 11, 12], provide robust frameworks for analyzing kinetic profiles and have been successfully applied to determine effective rate constants for various stages of aggregation. However, these rates emerge from multiple intermediate states of peptides and complex free energy landscapes, which cannot be completely reconstructed [1, 4]. Modern algorithms [13, 14] can predict aggregation-prone regions and structures of aligned fibril stacks, but elucidating oligomer structures remains challenging due to their elusive and transient nature [4]. Rate prediction is still underdeveloped, although kinetics are crucial for reparation systems and aging. Some models [10] successfully exploit the concept of the most unstable nucleus and the consequent minimal irreversibly growing seed. Such linear models can be modernized to elaborate on the underlying causes of the transition from instability to stability during oligomer growth. For example, spherical oligomer models could provide valuable insights into the nature of this instability, enhancing our understanding of the phenomenon [6]. One of the pioneering models describing the nucleation of a new phase [2], Zeldovich’s theory, has provided insights into the growth of protein crystals and the assembly of critical nuclei of peptide clusters [6, 7]. Furthermore, this model has been successfully applied to describe glass transition phenomena, particularly in the context of amorphous aggregation processes [3]. By comparing oligomer size distributions and kinetically estimated effective nuclei sizes, one can judge whether higher-order oligomers are involved in rearrangements toward protofibrils.

In this study, we examined the dependence of lag times on initial peptide concentrations and calculated the theoretical sizes of primary nuclei. Several theoretical kinetic regimes were explored. This approach yields insights into kinetic mechanisms from a single dataset, eliminating the need for comparisons across multiple sources with variable conditions, such as agitation levels, which significantly affect outcomes [1, 3]. Our findings align with critical micelle concentrations associated with the emergence of larger off-pathway oligomers, likely accounting for shifts in nucleation regimes observed at concentrations exceeding 10 µM for several peptides. We establish that a linear elongation model, even with the inclusion of fibril ends capping, is inadequate to account for the weak dependence of lag times on increasing concentrations. Incorporating secondary events, such as secondary nucleation or fragmentation, which produce additional growth sites, is critical for plausible modeling of peptide and protein aggregation.

## 2 Model of nucleation

We investigate oligomeric nucleation within the lag phase, which can be characterized in general as the interval during which disordered oligomeric species predominate over fibrillar assemblies. To elucidate this process, three mechanisms of primary nucleus formation have been proposed. In each mechanism, nuclei are characterized as spontaneously elongating ordered structures formed from conformationally reorganized disordered oligomers (Fig. 1) [6]:

**Fig. 1.**
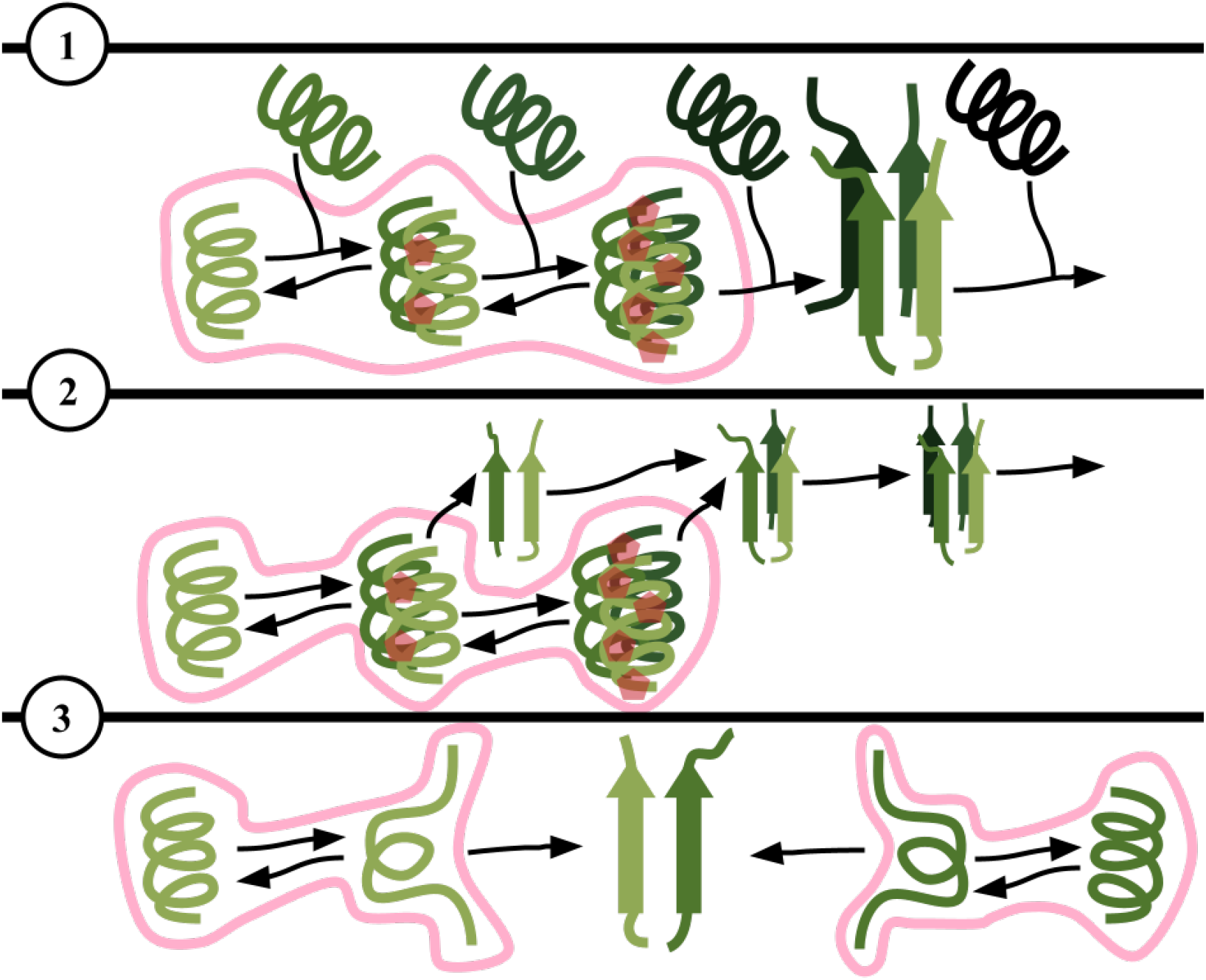
Diverse nucleation pathways. Most disordered oligomers dissociate back into monomers [6, 15, 16]. Red pentagons symbolize the transient interactions, primarily hydrogen and electrostatic bonds, and “tensions” within unstable oligomers. Pink curves represent species distributed approximately in accordance with the Boltzmann distribution during the lag phase.

**Fig. 2.**
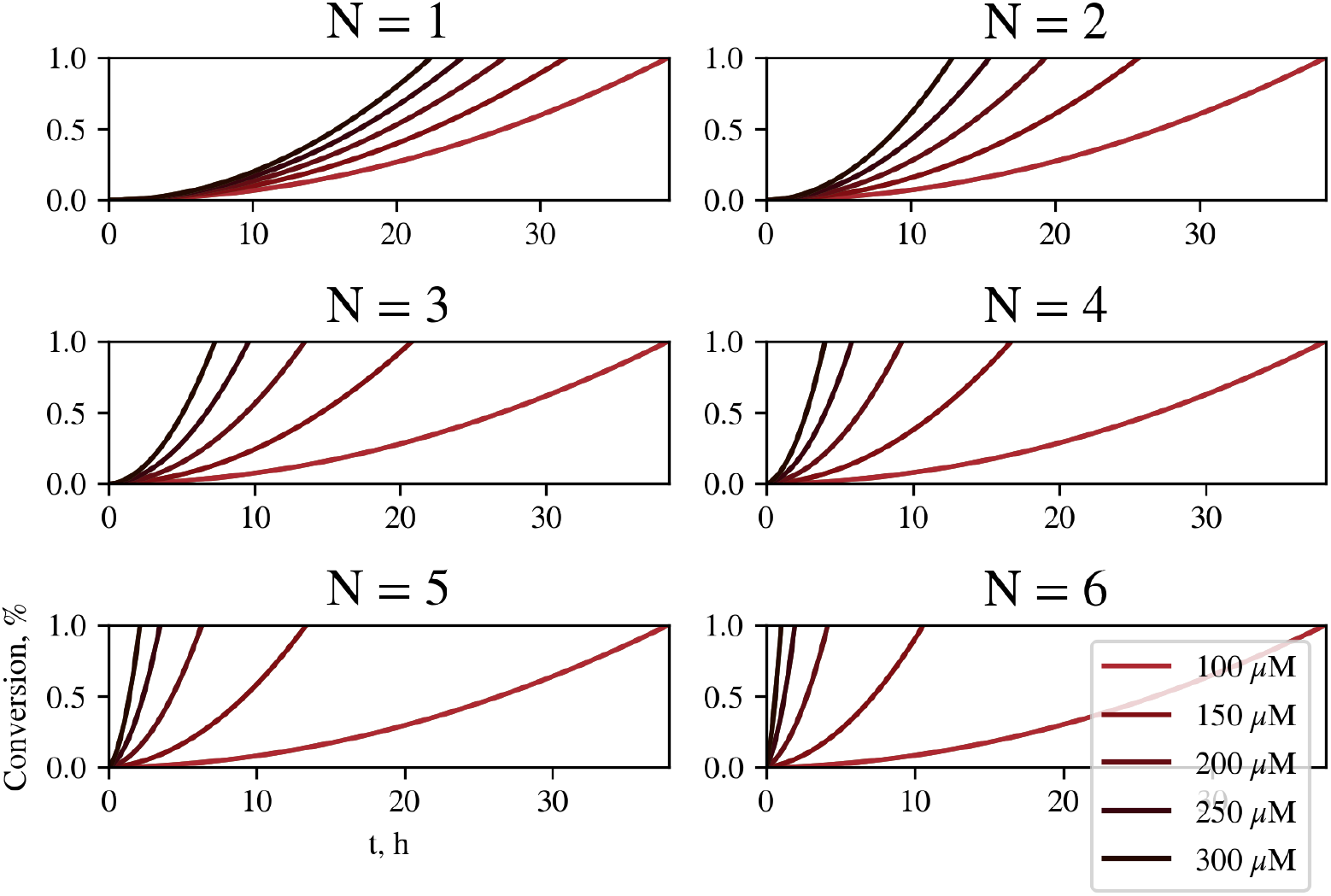
Theoretical kinetic curves during the lag phase under the elongation regime for varying primary nucleus sizes. Parameters differ only in *N* = 1, 2, 3, 4, 5, 6 and the corresponding *K*_*N*_ = 10^4(N −1)^ *M* ^1−N^.

**Fig. 3.**
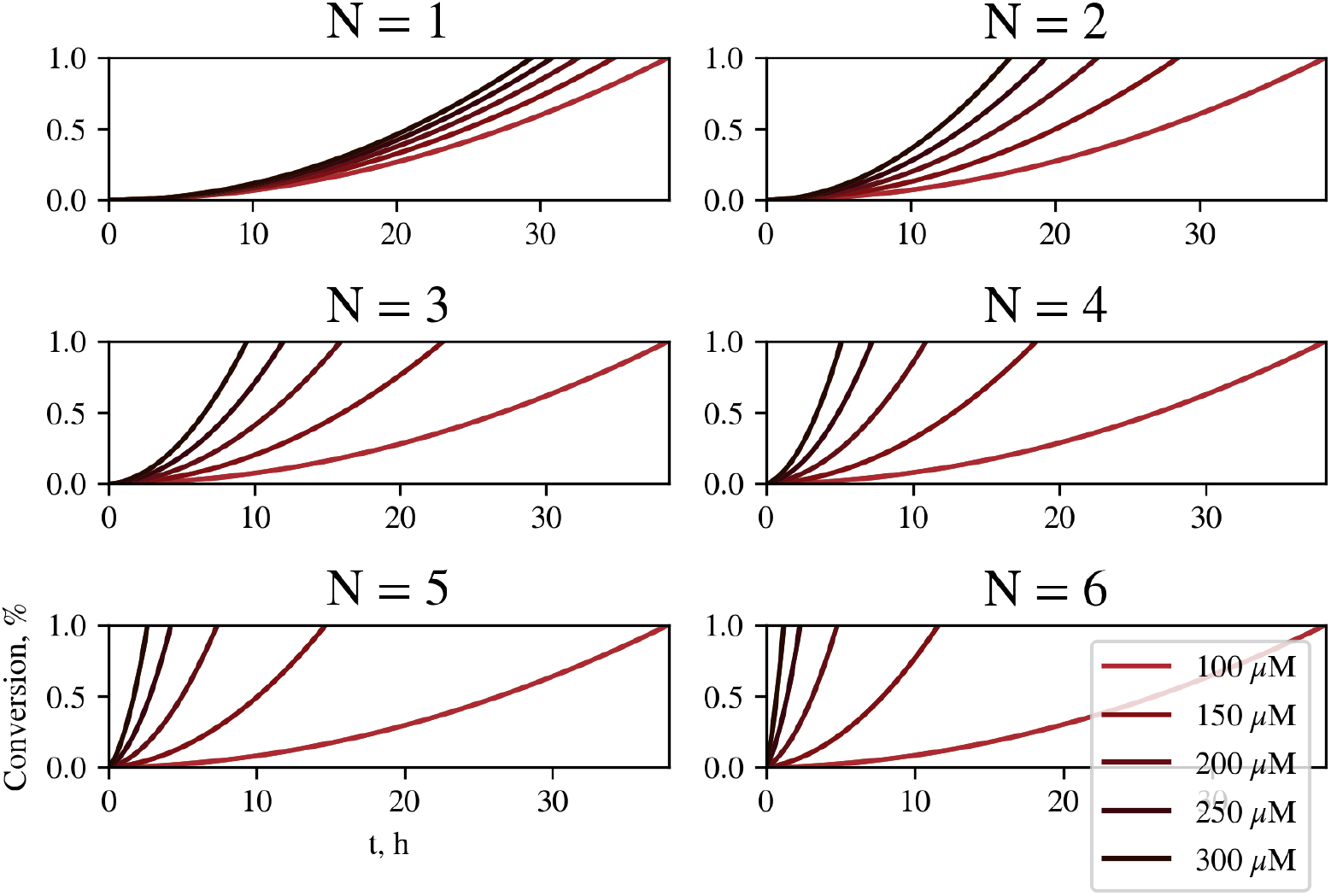
Theoretical kinetic curves during the lag phase under the elongation regime with capping (*c* = 2, *K*_cap_ = 10^4^ *M* ^−1^) for varying primary nucleus sizes. Parameters differ only in *N* = 1, 2, 3, 4, 5, 6 and the corresponding *K*_*N*_ = 10^4(*N*−1)^ *M* ^1−*N*^.

1. The nucleus can originate only from a prenucleus, defined as the most thermodynamically unstable oligomer that, upon addition of a single monomer, stabilizes and undergoes near-irreversible elongation [6, 15, 16].
2. Monomers rapidly assemble into disordered oligomers of varying sizes, each exhibiting distinct stability and propensity for conversion into ordered oligomers without changing their sizes, in monomers. The elongation of ordered oligomers enhances stability across all sizes, facilitating nucleation via multiple pathways as disordered oligomers transition into ordered nuclei. During the lag phase, disordered oligomers form a metastable distribution [3], progressively and irreversibly converting into nuclei [16], with conversion rates typically governed by oligomer size [3, 5, 6, 17].
3. Monomers in solution maintain a dynamic equilibrium between thermodynamically favored, (partially) folded conformations and less stable, (partially) denatured states. The nucleus, a dimer, forms with increased probability upon collision of two denatured monomers, while collisions of two native monomers are almost unproductive. This mechanism excludes nucleating oligomers larger than dimers, as ternary or higher-order collisions in solution are statistically negligible compared to binary interactions [3, 5, 18].

In approach (1), the prenucleus size *N* ^*^− 1 is defined as the number of monomers in an unstable oligomer that, upon addition of one more monomer, undergoes nearly irreversible growth. Consequently, the nucleation rate *r*, defined as the increase in nuclei concentration per unit time, is expressed as follows [2, 6, 8]:

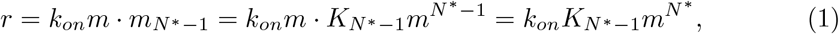

where *k*_*on*_ is the rate constant for monomer attachment to the prenucleus, *m* is the concentration of free monomers, 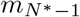 is the metastable concentration of prenuclear oligomers, and 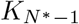 is the equilibrium constant for the formation of prenuclear (*N* ^*^ − 1)-mers. The parameters *k*_*on*_ and 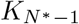 are averaged over diverse (*N* ^*^ − 1)-mer structures, which may exhibit considerable variability in their transition propensities. From (1) follows the expression for *N* ^*^, assuming that all peptides are initially free monomers, whose concentration does not decrease significantly during lag phase [6]:

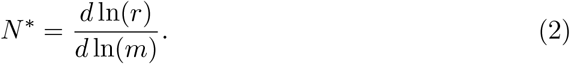

Unfortunately, direct measurements of the primary nucleation rate *r* are often challenging [9]. Instead, it is preferable to use a readily measurable quantity that is related to the nucleation rate. The lag time *t*_*lag*_ is one of the most accessible quantities to measure, as the lag phase is distinctly evident in kinetic curves [4, 19]. In protein aggregation studies, lag time is typically defined in two ways as the time: (1) at which the tangent to the maximum aggregation rate point intersects the time axis [8, 9], and (2) required for a critical fraction of molecules *α* to adopt fibril-prone conformations [4]. Reaching this critical fraction corresponds to the transition from a dynamic equilibrium between disordered oligomers to the rapid growth of fibrils [8]. The first method is compatible with various nucleation-growth models and facilitates analysis of growth-to-lag time ratios [10], but steep growth curves can lead to errors in estimating nucleus size. Experimental lag times from Fig. 10 were calculated in the cited studies using different definitions; however, for consistency, in our study we adopt only the second definition, assuming that the shift in the point of tangent intersection with the horizontal axis approximates the shift in the point of intersection between the curve and a selected horizontal line, i.e., both points are in the same small enough interval where significant growth initiates:

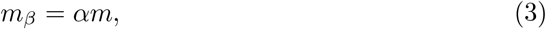

where *m*_*β*_ is the concentration of monomers converted into amyloids. One of the advantages of this definition is that it does not include the parameters of the growth and saturation phases, which may be accompanied by side transitions.

If secondary nucleation and fragmentation become significant already during the lag phase (heterogeneity [4, 10]), lag time and the rate of primary nucleation are no longer simply related as linear inverses. One of generalizations is power scaling in dimensionless units [8, 20]:

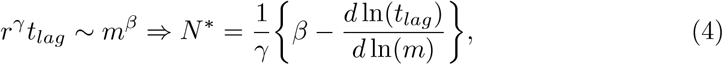

where *γ >* 0 is the rate scaling factor of primary nucleation arising from the influence of secondary nucleation and fragmentation, and *β >* 0 accounts for the increase in rate with concentration, reflecting that the switch from the lag to the growth phase requires other concentration of primary nuclei than a constant fraction, due to secondary events.

As usual, we suggest that fibrils elongate by monomer addition at both free ends with a rate constant 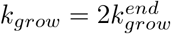 (we notice that fibrils with a single active plus-end 5 have been reported, but are not treated separately here; if required, the substitution 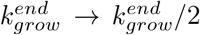 can be applied). Nucleation occurs via two mechanisms: primary nucleation from *N* -mers, governed by an equilibrium constant for transient *N* -mer association *K*_*N*_ and a kinetic rate constant for amyloid conversion *k*_*N*_, or secondary nucleation from *s*-mers adsorbed on fibril surfaces, characterized by a surface-assisted assembly constant *K*_*sec*_ and a conversion rate constant *k*_*sec*_. Fibrils can also break up into two new fibrils with a rate constant *k*_*frag*_. Although fragmentation is typically considered for seeds of size four or larger, we assume that even monomers resulting from dimer fragmentation retain an amyloid-prone conformation sufficiently long to enable monomer attachment before reverting to the native state [6, 15, 16]:

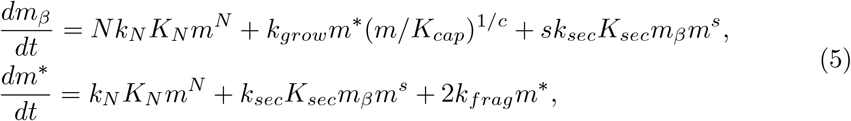

where *s* denotes size of the secondary nucleus, *c* is a size of capping oligomers which block fibril end, *K*_*cap*_ is an equilibrium constant of capping; if capping events are negligible, *c* = 1 and *K*_*cap*_ = 1, *m*^*^(*t*) is the concentration of growing amyloid aggregate active ends at time *t*, and the critical fraction (3) of converted monomers should be selected of order *α ∼ k*_*olig*_*/k*_*fibr*_, where *k*_*olig*_ and *k*_*fibr*_ are the mean rate constants of monomer attachment to an oligomer and to a fibril, respectively. If secondary nucleation is taken into account, *α ∼ k*_*olig*_*/k*_*sec*_ [9]. Fortunately, switching to the derivatives in double logarithmic scale removes *α* from consideration. For convenience, we introduce one-letter notation for kinetic coefficients in (5):

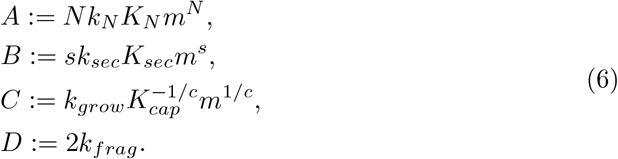

and assume that *m*(*t*) *≈ m*(0) during the lag phase, which is usually valid for *α ≪* 1. We now estimate the error associated with this approximation. If the derivative of a function *y* can be expressed as a polynomial of the variable *x* with coefficients {*a*_*p*_}, and an approximate value *x*_*a*_ is used instead of its true value, an upper bound for the relative rate error can be derived:

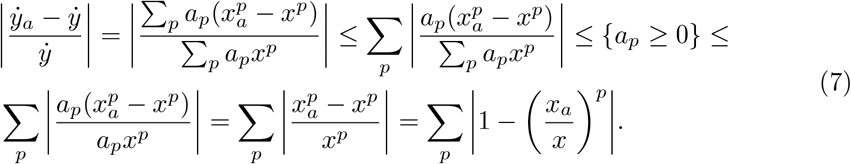

We note that the kinetic equations (5) contain no more than three such terms. Furthermore, our analysis considers exponents within the bounds *N* ≤ 6, *c*^−1^ ≤ 1, and *s* ≤ 2. Under these constraints, the relative error in the primary nucleation rate does not exceed 9%. In practice, the exponent *N* is typically ≤ 4, which reduces the error to approximately 7%. However, as demonstrated in the subsequent analysis, the error is even smaller. The actual uncertainty is further reduced because the concentration of free monomer enters the rate equations for secondary processes with lower powers, and the contribution from secondary pathways typically dominates that of primary nucleation.

The precise solution for the aggregated mass *m*_*β*_ in (5) is expressed through the eigenvalues *λ*_1_ and *λ*_2_ of corresponding characteristic matrix:

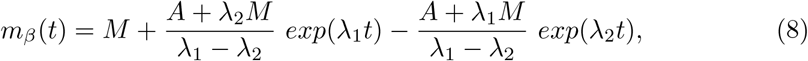

where *M* is the partial solution of heterogeneous system of linear ODEs (5):

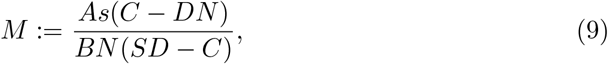

and the eigenvalues of the characteristic matrix of (5) are the solutions of the corresponding quadric equation, as the system contains only two variables:

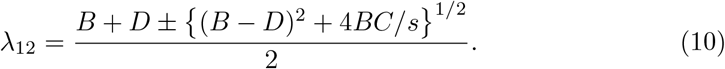

When fragmentation is neglected, the exact solution of (8) can be approximated by a quadratic polynomial. The simplified expression:

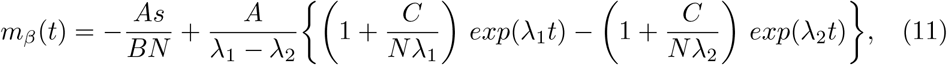

the eigenvalues in which emerge as roots of the characteristic quadratic equation governing the dynamics in (11) if *D* = 0:

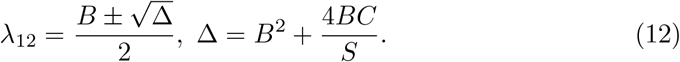

Under the assumption that *λ*_1_*t, λ*_2_*t ≪* 1 during the lag phase, we approximate *m*_*β*_(*t*) with a polynomial of *t*, yielding a second-order Maclaurin expansion for *m*_*β*_(*t*).

This simplifies when *C ≫ B*, indicating that elongation dominates over secondary nucleation in the lag phase:

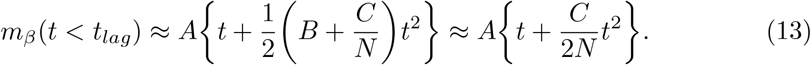

That means, the lag time calculation for a specified concentration is reduced to a quadratic equation:

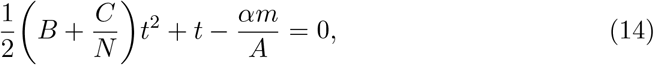

which has exactly one non-negative solution because all parameters are non-negative and *C >* 0:

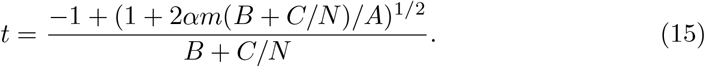

To elucidate the concentration dependence of the solution, we explicitly express the powers of *m* in the kinetic coefficients, introducing new coefficients that are independent of *m*, either as constants or as functions of *N* :

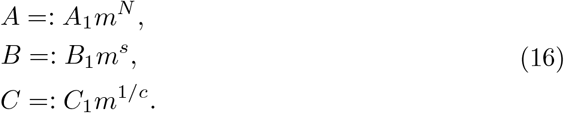

Using this notation, (15) can be reformulated in terms of powers of *m* and transformed to a double logarithmic scale:

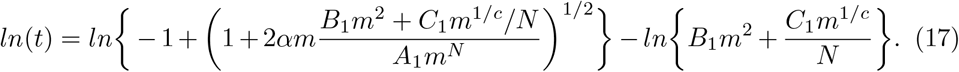

From (17), the slope of the logarithmic plot can be derived using the relation *d* \**/d* ln(*m*) = *m* (*d*\* */dm*) in the right-hand side, where represents any smooth enough function:

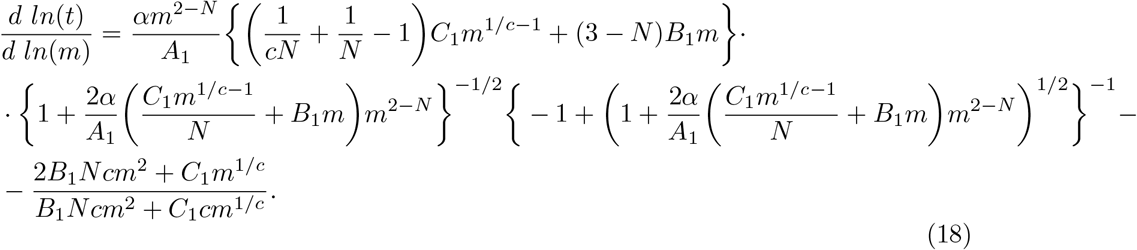

When *B*_1_*m ≪ C*_1_*/N* (⇔ *B ≪ C/N*), two limiting cases can be evaluated based on *A*_1_, employing the appropriate Maclaurin expansion (1 + *x*)^1*/*2^ *≈* 1 + *x/*2 for *x ≪* 1, and (1 + *x*)^1*/*2^ *≈ x*^1*/*2^ for *x ≫* 1:

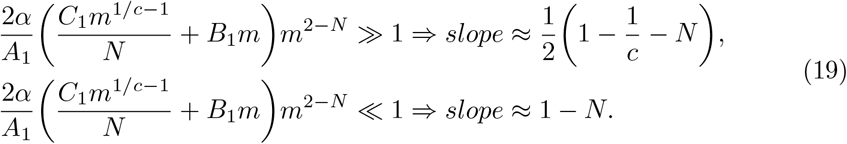

Specifically, from (19), it follows that for *N* ≥ 2, the absolute slope satisfies |*slope*| ≥ 1 in the elongation regime and |*slope*| ≥ 0.75 under elongation with capping. This derivation indicates that the lag time is not simply the inverse of the nucleation rate. Assuming that |*slope*| is small enough, from (19), the primary nucleus size can be expressed in terms of the lag time *t*_*lag*_ [8] as the maximum of two approximations:

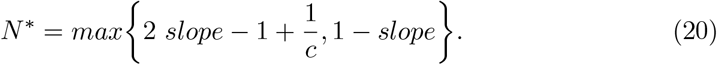

However, when *c* = 1, no combination of parameters *A*_1_, *B*_1_, *C*_1_ can account for slopes less than unity [21] (Table 1). This suggests that competitive capping of fibril ends may occur, with capping oligomers occupying growing ends while concentrations remain relatively high during the lag phase.

**Table 1.**
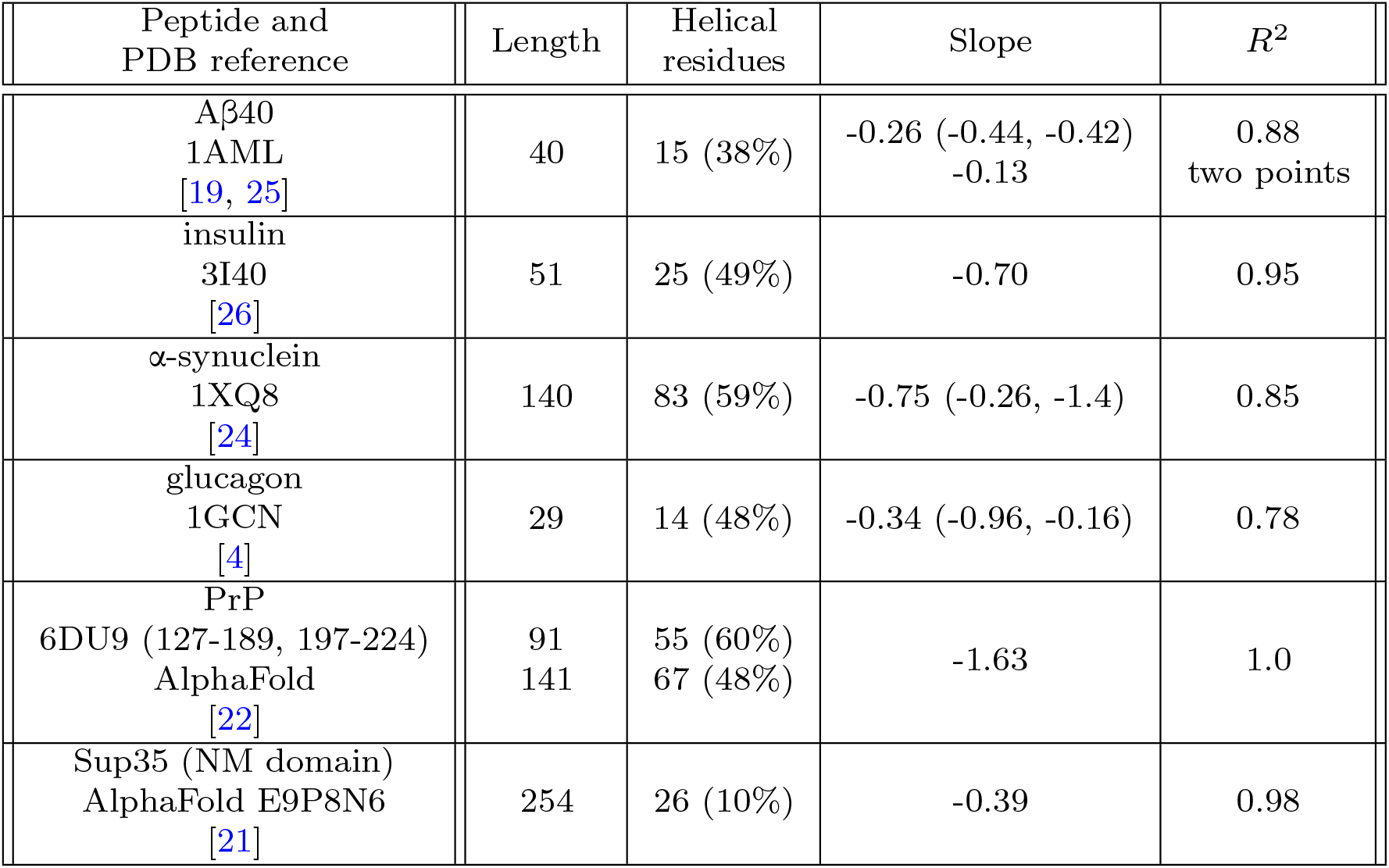
Secondary structures of peptides determined by DSSP, and nucleation parameters derived from the double logarithmic plot in Fig. 10. For some peptides, separate estimations for *m* below and above possible regime switch concentrations are provided in brackets.

| Peptide and PDB reference | Length | Helical residues | Slope | $R^2$ |
| --- | --- | --- | --- | --- |
| Aβ40<br>1AML<br>[19, 25] | 40 | 15 (38%) | -0.26 (-0.44, -0.42)<br>-0.13 | 0.88<br>two points |
| insulin<br>3I40<br>[26] | 51 | 25 (49%) | -0.70 | 0.95 |
| α-synuclein<br>1XQ8<br>[24] | 140 | 83 (59%) | -0.75 (-0.26, -1.4) | 0.85 |
| glucagon<br>1GCN<br>[4] | 29 | 14 (48%) | -0.34 (-0.96, -0.16) | 0.78 |
| PrP<br>6DU9 (127-189, 197-224)<br>AlphaFold<br>[22] | 91<br>141 | 55 (60%)<br>67 (48%) | -1.63 | 1.0 |
| Sup35 (NM domain)<br>AlphaFold E9P8N6<br>[21] | 254 | 26 (10%) | -0.39 | 0.98 |

**Table 2.**
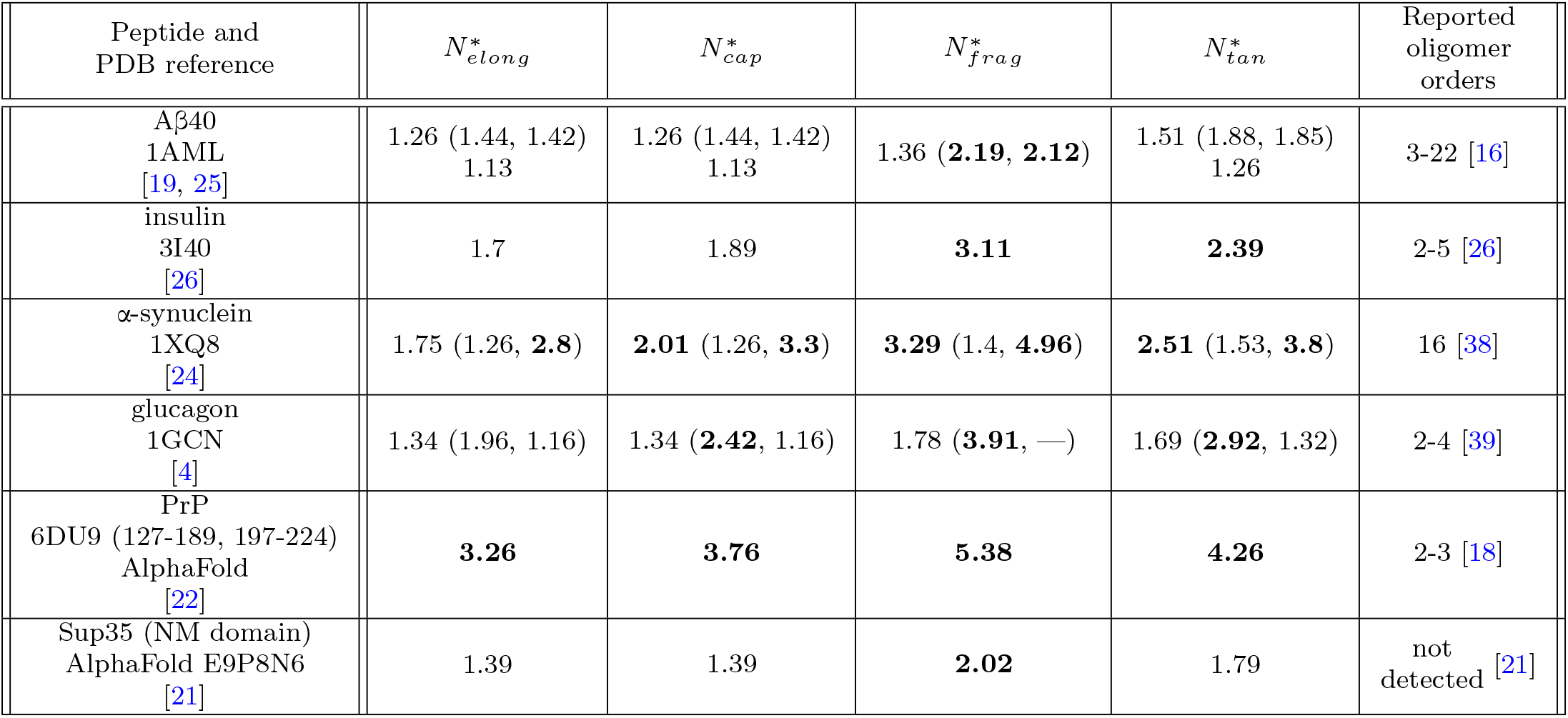
Critical nucleus sizes 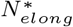 and 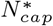 were calculated using equation (20) with parameter values *c* = 1, *K*_*cap*_ = 1 and *c* = 2, *K*_*cap*_ = 10^4^, respectively. Fragmentation-regime theoretical sizes have been calculated using interpolation method (21, 22) with pre-calculated slope table for nuclei sizes: 1 → −0.18, 2 → −0.39, 3→ − 0.66, 4 → − 0.99, 5→ − 1.42, 6 → −1.97. For some peptides, separate estimations for *m* below and above possible regime switch concentrations are provided in brackets. Theoretical nuclei sizes equal or greater than two are highlighted bold and assumed as acceptable.

From our computations, it is evident that in the fragmentation-driven model, the quantities *λ*_1_*t*_*lag*_ and *λ*_2_*t*_*lag*_ are not sufficiently small to permit the application of Maclaurin expansion, which would simplify the derivation of the theoretical slope to a quadratic equation. To address this, we pre-computed theoretical tables of approximate expected slopes for specified kinetic constants. Using these tables and an observed experimental slope, one can estimate *N* through the momentum-rule approximation:

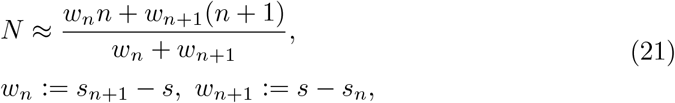

where [*n, n* + 1] is an interval containing *N* if |*s*(*n*) | is a monotonically increasing function, i.e. *n* must satisfy the following condition:

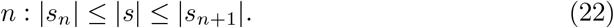

The linear dependence of *ln*(*t*_*lag*_) on *ln*(*m*) has been observed across various peptides [10]. For instance, in first-order transitions such as prion misfolding, where peptides irreversibly adopt a pathological conformation with *d ln*(*t*_*lag*_) */d ln*(*m*) = 0, monomeric nucleation is expected, yielding *N* ^*^ = 1. However, such stable prion-like states are rare among peptides, necessitating the inclusion of secondary processes like fragmentation. Consequently, we incorporate fragmentation even during the lag phase, as its omission bans the theoretical framework from accounting for many experimentally observed slopes. Amyloid propagation driven by fragmentation alters the functional dependence from linear to concave (Fig. 4).

**Fig. 4.**
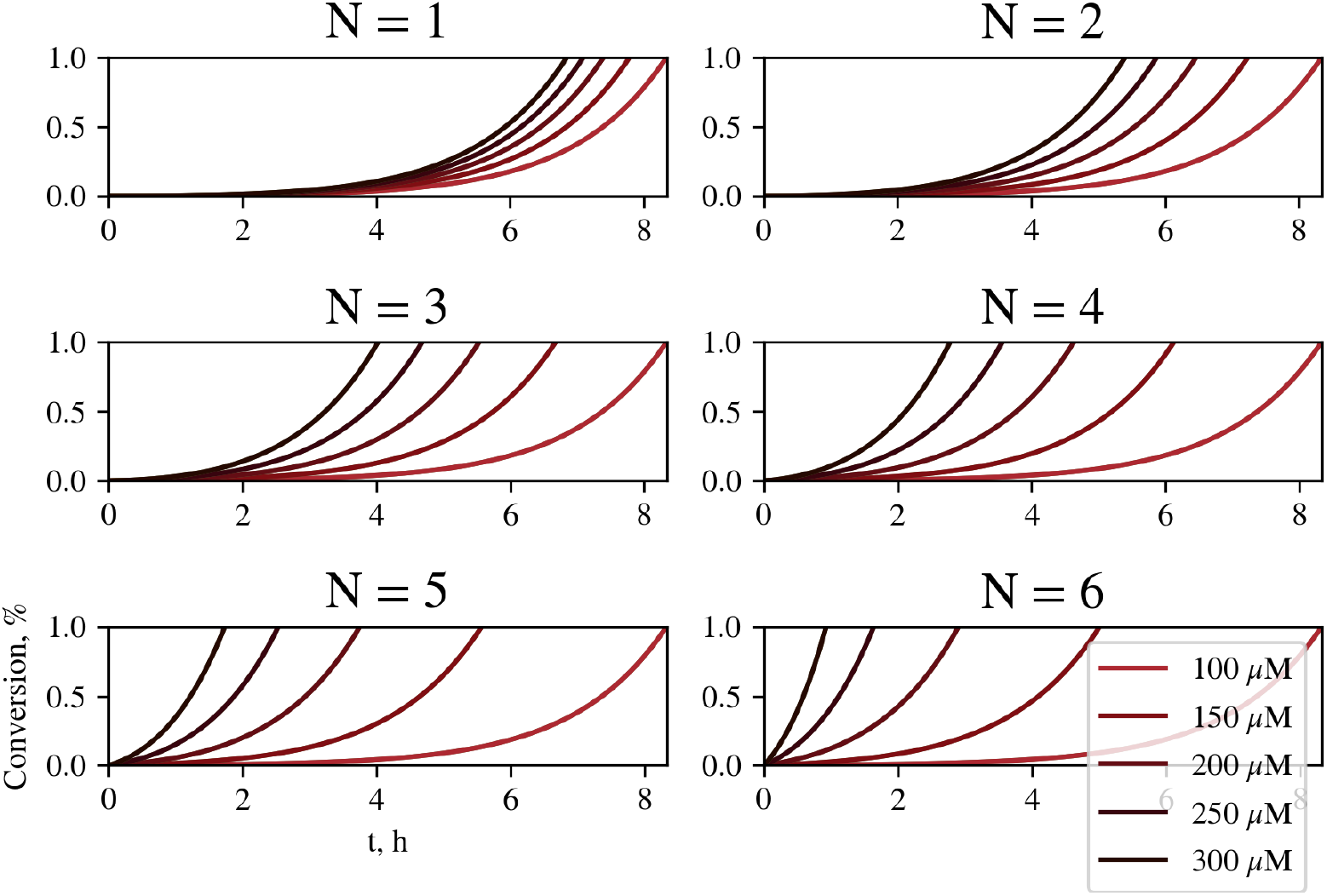
Theoretical kinetic curves during the lag phase under the fragmentation regime (*k*_*frag*_ = 10^−4^ *s*^−1^) with capping (*c* = 2, *K*_cap_ = 10^4^ *M* ^−1^) for varying primary nucleus sizes. Parameters differ only in *N* = 1, 2, 3, 4, 5, 6 and the corresponding *K*_*N*_ = 10^4(*N*−1)^ *M* ^1−*N*^.

**Fig. 5.**
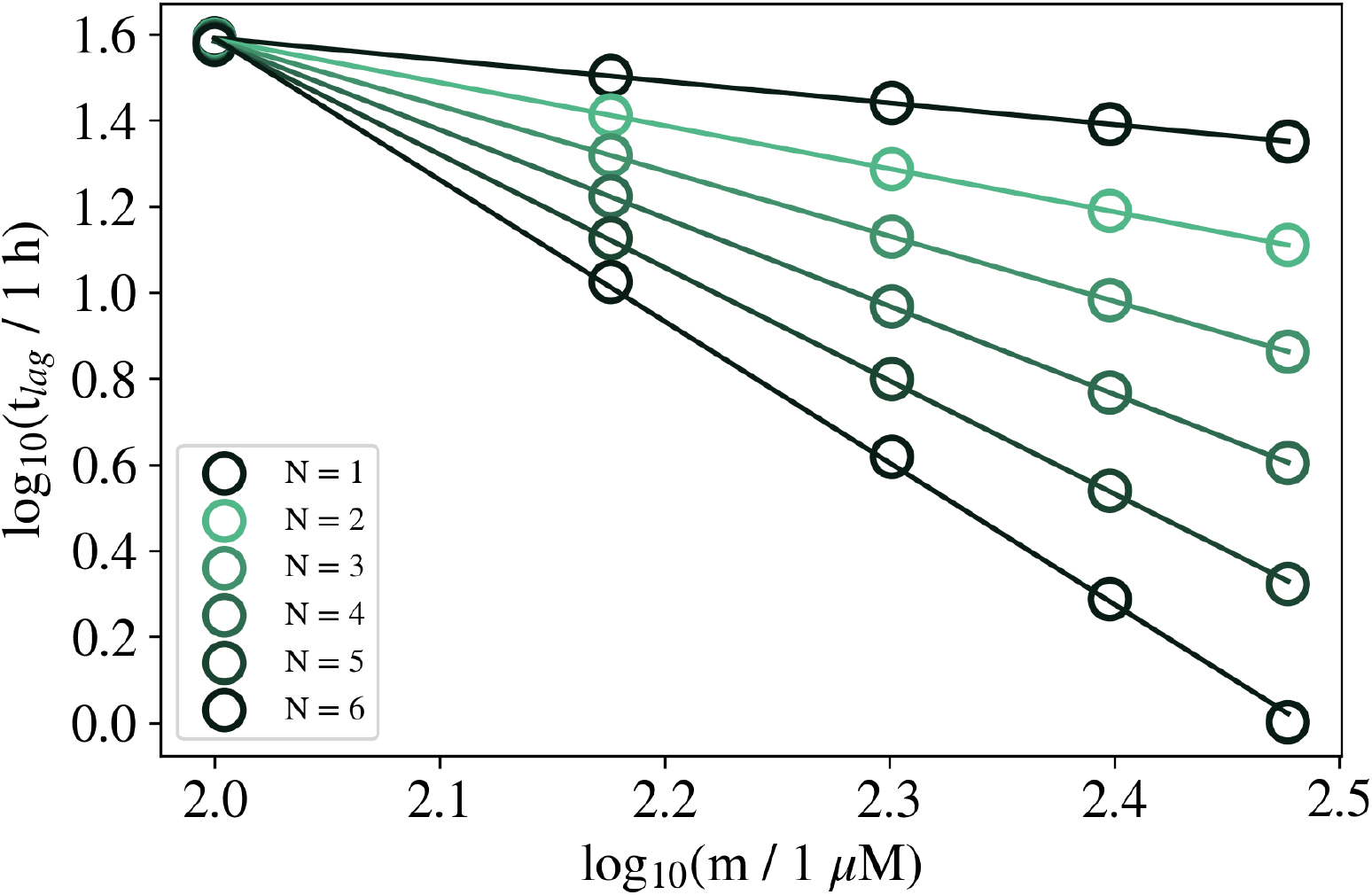
The theoretical concentration dependencies of lag times calculated for nucleus sizes ranging from 1 to 6 under the elongation regime.

**Fig. 6.**
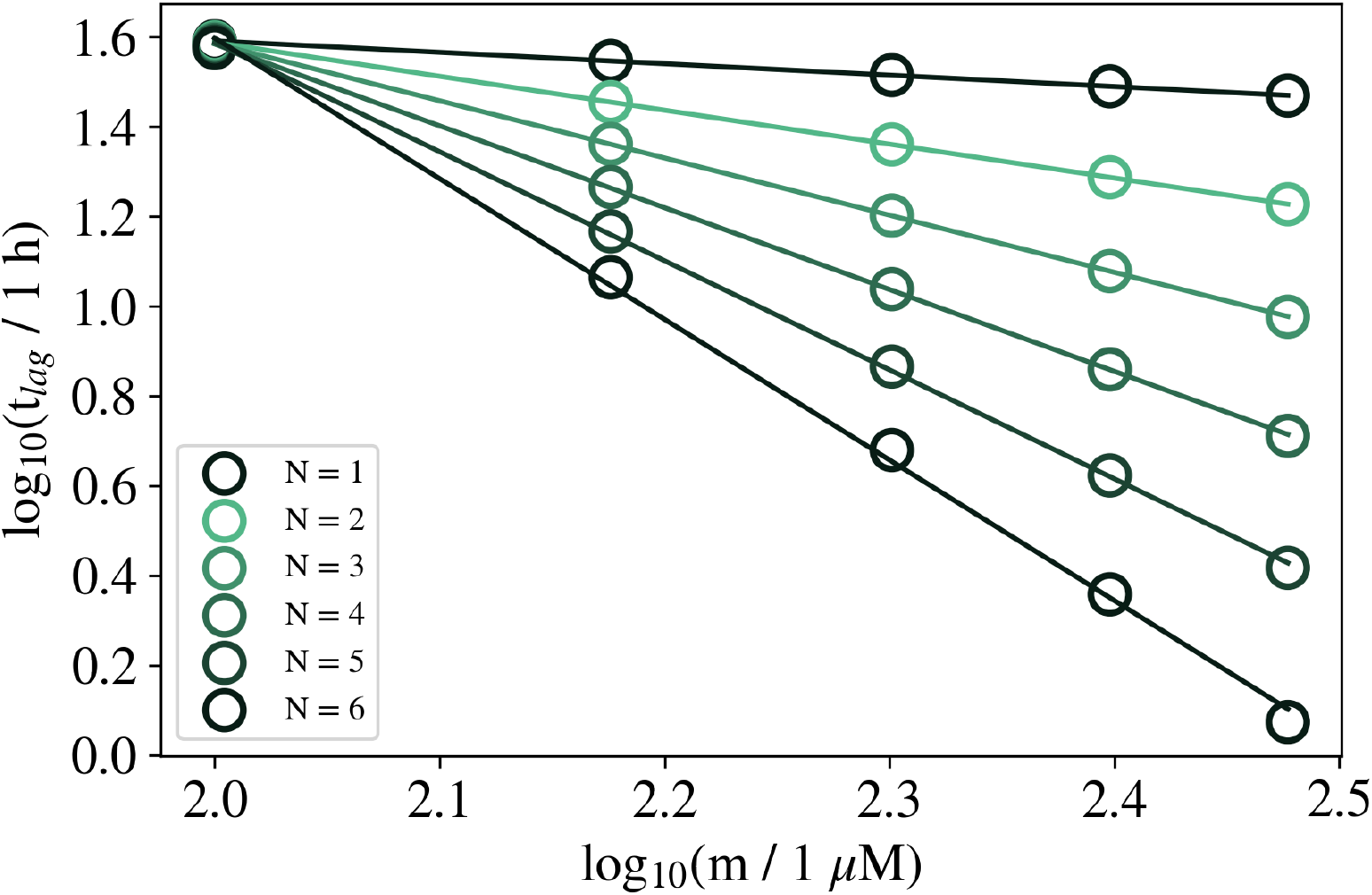
The theoretical concentration dependencies of lag times calculated for nucleus sizes ranging from 1 to 6 under the elongation regime with capping.

**Fig. 7.**
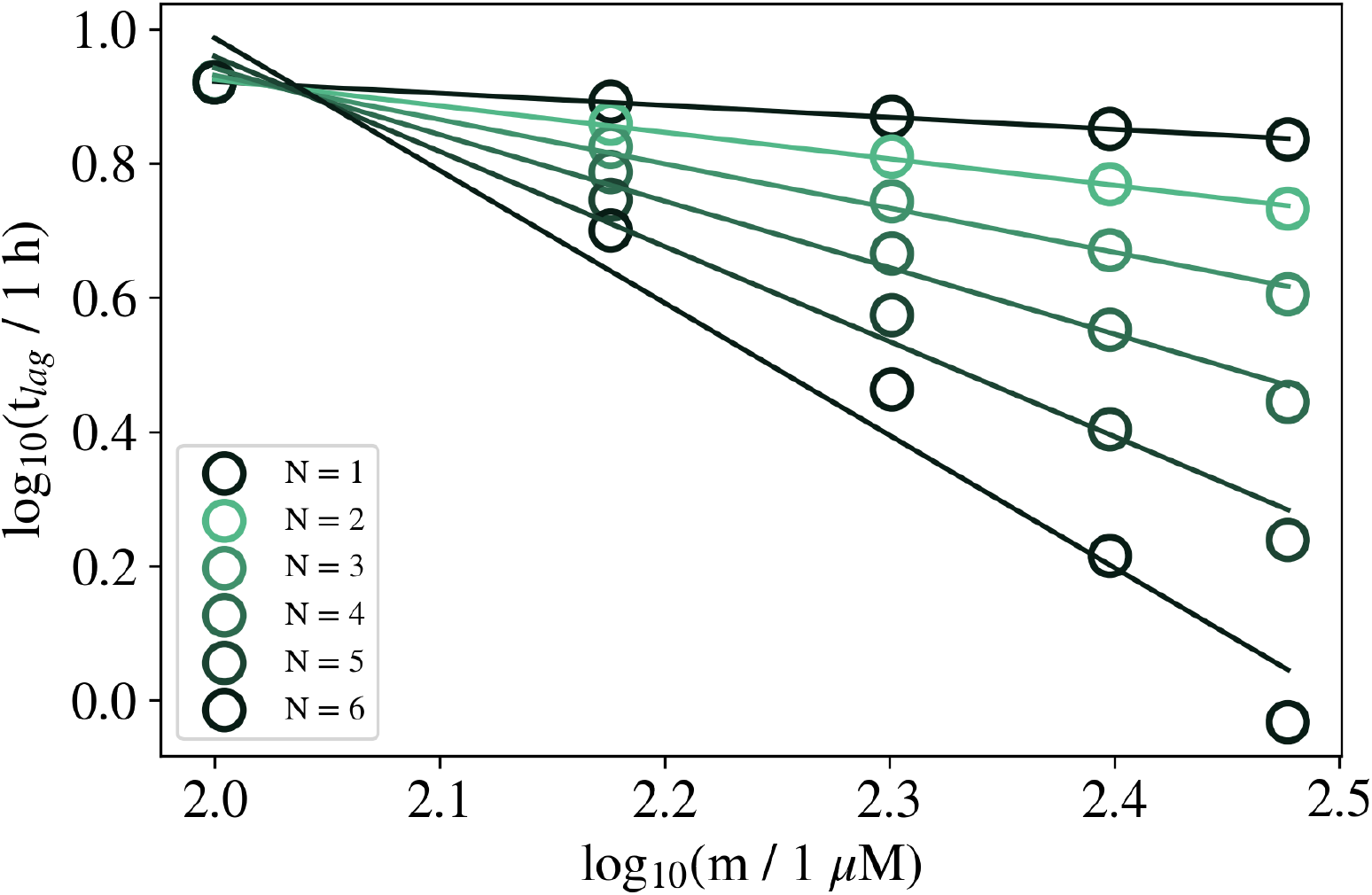
The theoretical concentration dependencies of lag times calculated for nucleus sizes ranging from 1 to 6 under the fragmentation regime.

**Fig. 8.**
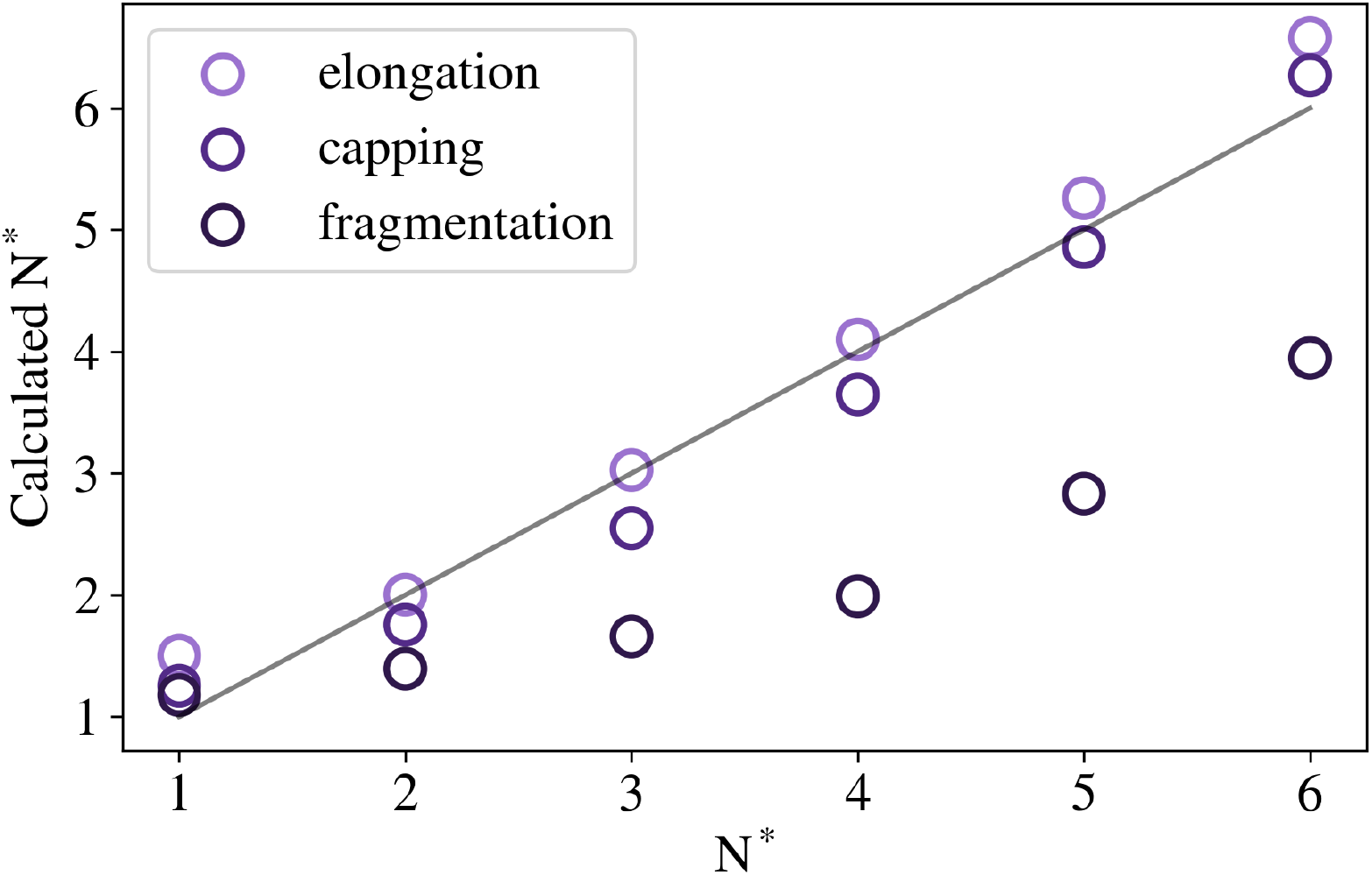
Primary nuclei sizes used in the model calculations and their predictions using (20). Gray line denotes ideal coincidence.

**Fig. 9.**
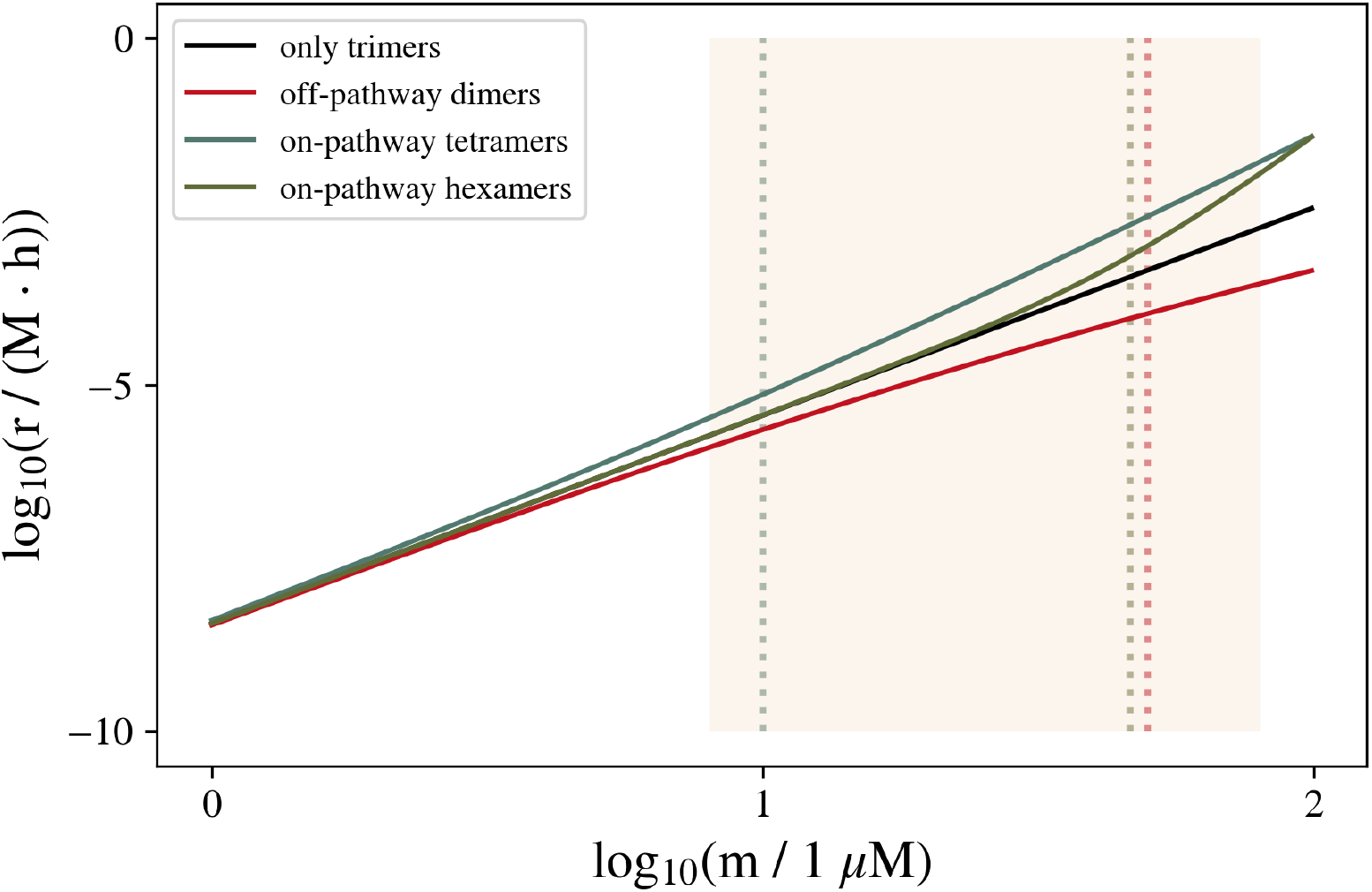
Theoretical dependencies of the primary nucleation rate on concentrations, accounting for multiple on- and off-pathway oligomer sizes.

At the same time, using the definition of the lag phase based on the tangent to the highest-rate point of the growth phase, the duration of the lag phase for linearly growing polymers has been derived [10]:

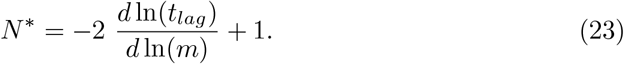

Although (23) and (20) give distinct values of *N* ^*^, both order peptides by nucleus size in the same way, larger slopes correspond to larger nuclei. We remind the reader that we define the nucleus as the smallest stable and converted oligomer, rather than the largest unstable one, as some authors do [6, 10]. This discrepancy can cause differences of *±*1 between *N* ^*^ estimations, which should be attributed to differing definitions rather than differences in the models.

For all model calculations, the following parameters have been used: *α* = 0.01, *s* = 2, *k*_N_ = 10^−9^ *s*^−1^, *K*_N_ = 10^4(N −1)^ *M* ^1−N^, *k*_sec_ = 10^−4^ *s*^−1^, *K*_sec_ = 10^3^ *M* ^−2^, *k*_*grow*_ = 10 *M* ^−1^*s*^−1^, *t*_*max*_ = 40 *h*.

The nucleation rate is more precisely described by the Zeldovich factor [6, 7, 15], which quantifies the probability that an unstable oligomer binds the final monomer before dissociating:

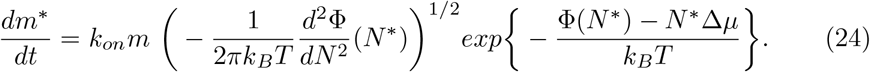

This equation links the kinetics of nucleation with its free energy profile. Here, Φ(*N* ^*^) is the free energy of an *N* -mer relative to *N* free solvated monomers under standard conditions, and Δ*µ* = *k*_*B*_*T ln*(*m/m*_*sat*_) [7] represents the change in monomer chemical potential, derived under the ideal solution approximation applicable for sufficiently dilute samples, while typical experimental concentrations *m ∼* (10 *÷* 1000) µM. In this expression, *k*_*B*_ is the Boltzmann constant and *T* is the absolute temperature, typically ranging from 20°C to 60°C.

However, this equation is difficult to apply to peptide oligomerization because it was originally developed for sufficiently large formations such as bubbles, whereas peptide nucleation is usually driven by small oligomers of 1 to 4 monomers (Table 1). Moreover, peptide oligomerization lacks a regular free energy function analagous to those for growing bubbles and droplets, which are described by contributions of bulk phase transition and surface tension [7]. This complicates the interplay between the attachment rate constant and the shape of the free energy profile near the nucleus maximum. Additionally, this model does not account for multiple nucleation pathways with differing critical free energies; a few oligomers following a lower-barrier pathway can be far more productive for seeding than millions of seeds on side pathways. Finally, the model assumes a straightforward definition of free energy based solely on the number of monomers and initially relies on the ratio between surface tension and bulk contributions. This assumption does not hold for small clusters of macromolecules, where internal transitions and fluctuations play a more significant role.

If one neglects the decrease in *m* during the lag phase and assumes that equilibrium concentrations of oligomers are established much faster than their conversion [22], the nucleation pathways can be summed to obtain the resulting concentration of nuclei after a given time from the start of the experiment:

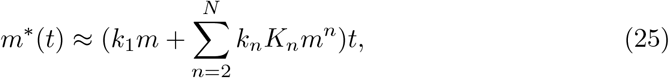

where *m*^*^(*t*) is the concentration of nuclei, *k*_1_ is the rate constant for monomer conversion to an aggregation-prone conformation, *k*_*n*_ are the rate constants for *n*-mer conversion, *K*_*n*_ are the equilibrium constants for the assembly of *n*-mers, and *t* is the time since peptide solvation. Sum (25) gives one of explanations why the effective nucleus size (Table 1) could be far from integer: it is a sum of contributions from different nucleation pathways involving oligomers of various sizes. However, we cannot exclude also the possibility of multistage mechanisms involving partial denaturation and induction, which could also produce non-integer effective reaction orders for nucleation. For example, assuming that nucleation starts from two adjacent peptides within an oligomer, while their surroundings act as a facilitating matrix, the slope can increase with concentration due to a larger fraction of higher-order oligomers.

For a pair of primary nucleation mechanisms, the critical concentration to switch the main nucleus size follows from (25):

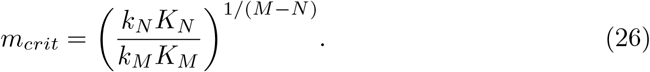

In contrast to serving as nucleation matrices, large oligomers may also act as peptide accumulators, slowing their β-conversion and reducing the nucleation rate as concentration increases. For instance, if *N* -mers are the main productive species while *N*_*off*_ -mers serve as depots [1] prevailing at high concentrations, above which *m_off_ N_off_ ∼ m*:

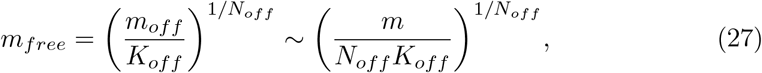

where *m*_*free*_ denotes the concentration of free monomers that are not involved in off-pathway structures and remain available for nucleation. Off-pathway species always reduce the concentration of free monomers.

In this case, the lag time is expressed in terms of the low concentration of free monomers (27):

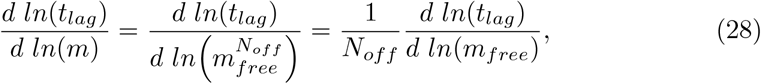

leading to a modified expression for the effective nucleus size for regimes where *N ∝ slope*:

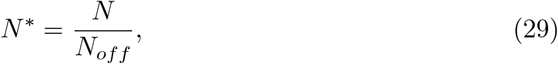

which means that off-pathway oligomers can reduce *N* ^*^ to values even less than one monomer if *N*_*off*_ *> N*. Since *N*_*off*_ ≥ 2, obtaining *N* ^*^ *>* 1 under high concentrations in the presence of multiple large oligomers provides evidence for matrix-assisted nucleation.

Within the framework of models (25) and (28), the estimated change in the nucleation rate slope corresponds to the critical micelle concentration (CMC), expressed either in fractional units (*c*) [23] or, as used here, in molar units:

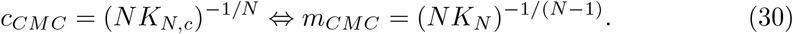

The equilibrium constants of oligomerization are calculated as 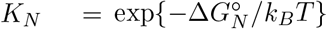. The partial free energy change of a monomer upon transition into an *N* -mer is 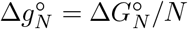. Such CMC estimates are applicable to both on-pathway and off-pathway aggregation models. On-pathway higher-order oligomers result in a steeper negative slope in concentration-rate dependence, as more species contribute to shortening the lag time. Conversely, off-pathway oligomers lead to a less negative slope, since additional assemblies slow down the conversion process.

## 3 Results and discussion

To estimate the effective nucleus size from lag time data, we analyzed the experimental points, initial monomer concentrations *m* and lag times *t*_*lag*_, collected from the studies referenced in the description of Fig. 10 and Table 1. These data were plotted on a double logarithmic scale with attempt to linearize the relationship between *m* and *t*_*lag*_ (Fig. 10). For each series we performed a linear least-squares regression to determine the slope of the best-fit line. The slopes obtained from these regressions are summarized in Table 1. The slopes of the regression lines range among typical orders 0.1 *÷* 1.0 and correspond, within the applied model, to the effective nuclei sizes of *N* ^*^ *≈* 2 *÷* 4. Small nucleus sizes up to 5 have also been reported in [10], based on the duration of the growth phase and its ratio to the lag time. That model incorporates secondary nucleation at the fibril surface, fragmentation, and bifurcation. Such values of *N* ^*^ provide evidence against the applicability of (24) to primary nucleation of peptide aggregates. Indeed, for a small nucleus size of order *N* ^*^ *≈* 2, the corresponding prenucleus is almost monomeric, which undermines the relevance of the Zeldovich convexity factor, assuming the existence of an interval in phase space around *N* ^*^. Therefore, instead of continuous pathways, a discrete cluster enumeration approach appears to be a more appropriate description. Such approach is widely used in molecular dynamics studies [6, 31]. These typical for protein aggregation nucleus sizes are consistent with estimates obtained by other methods. For example, the concentration dependence of another kinetic metric of aggregation, the time of semi-saturation *t*_50_, depending significantly on agitation level of a sample [1], demonstrated for Aβ42 slopes − (0.62 *÷* 1.33), which correspond again to *N* ^*^ *≈* 2 *÷* 4 [20]. It is intriguing that under the highest agitation level the authors almost reached the theoretical limit of slope corresponding to pure fragmentation, which indicates switch to another mechanism.

**Fig. 10.**
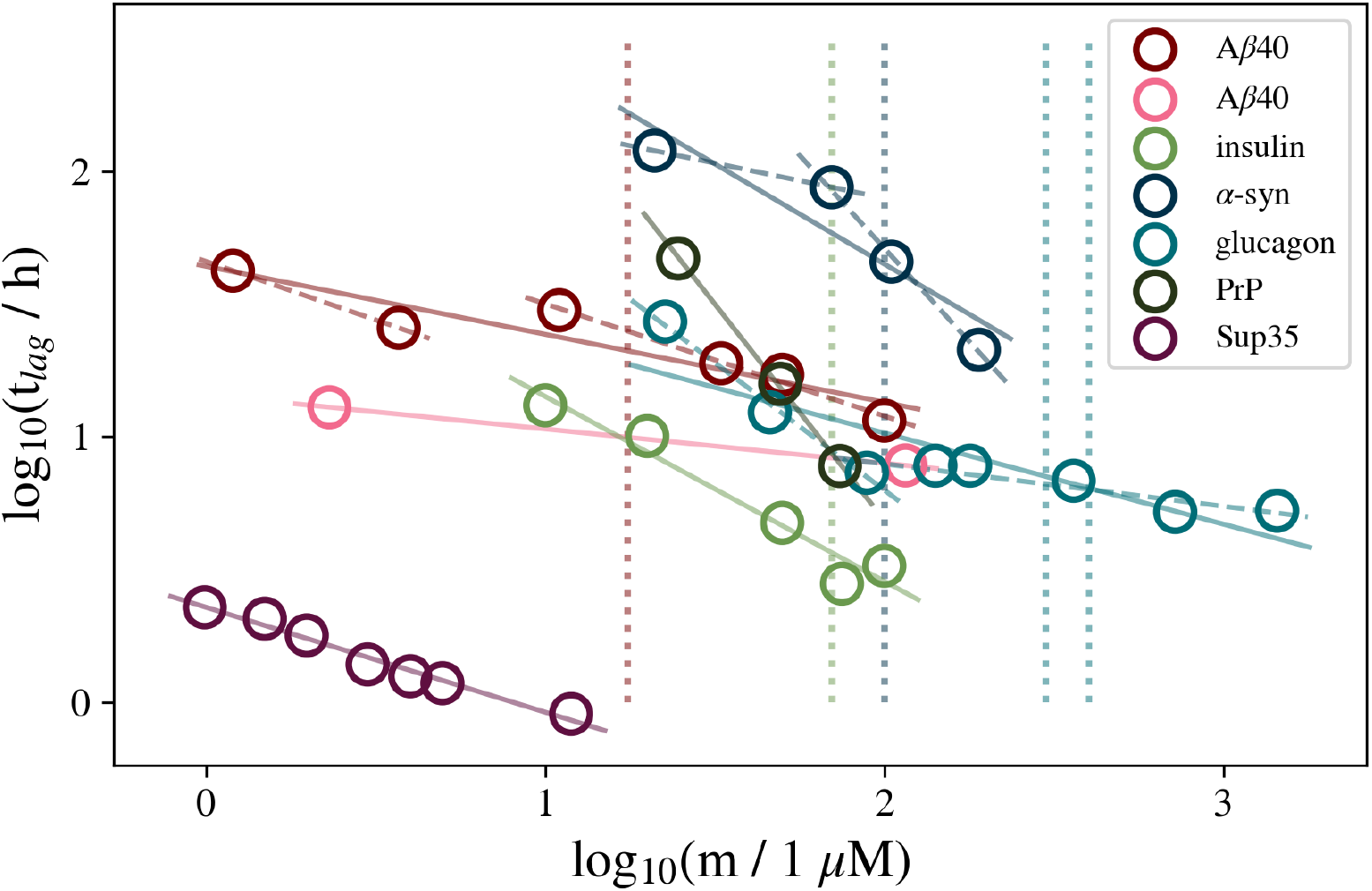
Aggregation lag times versus initial concentrations of different peptides. Data sources are experimental studies on α-synuclein ([24]), Aβ40 ([25], [19]), insulin ([26]), glucagon ([4]), PrP ([22]), Sup35 (NM domain) ([21]). The blue vertical line indicates 100 µM, approximately the concentration above which α-synuclein micelles likely convert to oligomers [27]. The red line marks the CMC of Aβ40 [28], the green line corresponds to the CMC of insulin hexamers [29], and the vertical blue stripe covers the approximate range of CMCs for glucagon [30]. Colored dashed lines represent separate regressions for the presumed subcritical and supercritical regimes. Circle sizes do not represent errors.

Lower concentrations yield higher linearity, with the lowest *R*^2^ value obtained for glucagon presenting the highest concentrations in its series and one of the lowest *R*^2^ appeared for Sup35 under the lowest concentration range. It aligns with the CMC model explaining switch from lower-order regimes (equation (1)) to higher-order regimes (equations (25) and (28)), resulting in a changing slope of the {*log*(*m*), *log*(*t*_*lag*_)} lines. However, while concentration decreases, usually not only the lag time, but also its and its standard deviation increases, meaning that more measurements may be required to construct the same confidence interval [19, 32]. Since most of the mentioned CMCs are greater than 10 µM, conducting kinetic experiments at concentrations *m <* 10 *µM* might be the preferred strategy for determining nucleus size, especially keeping in mind that physiological concentrations are usually even lower [1]. One possible explanation for why two selected subseries from the same Aβ40 dataset are nearly parallel is that the nucleus in higher-order oligomers may remain dimeric [6], with minimal influence from surrounding peptides. In this case, shifting the equilibrium from dimers to trimers, tetramers, and so on, has little effect on the nucleation pathway, but can slow the process resulting in a vertical translation of the line, as seen in Fig. 10. Although the entire series for Aβ40 could be interpreted as a single linear dependence with measurement errors, the explanation involving two lines has the advantage of accounting for the CMC.

At concentrations in range 10 *÷* 100 µM, several peptides demonstrate a change in the slope, which, in the context of the CMC model, corresponds to 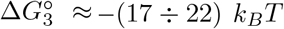 or 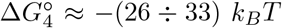. This is equivalent to standard average oligomerization free energy per monomer of Δ*g*° *≈* −(6 *÷* 8) *k*_*B*_*T*, and at 100 *µ*M the free energy of Δ*g*(10 ÷100 *µ*M) ≈ (3 ÷4) *k*_*B*_*T*, which are consistent with free energy profiles calculated using molecular dynamics simulations [17, 33]. One of peptides with possible slope change is glucagon; this hormone is known for its ability to associate into dimers, trimers, and hexamers, and corresponding free energies of association obtained from optical rotation measurements are 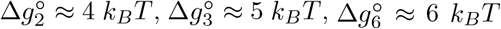, [30], corresponding to CMC range (300 *÷* 400) µM. The discrepancy in CMCs between different studies may be attributed to variations in pH [1] in [4] and [30], as well as to the presence of negative ions in the buffer used in [4], which may shield positive residues and facilitate aggregation [1]. In contrast, lag times for PrP and Sup35 at lower concentrations show strong linear dependencies, suggesting that these systems remain below their respective CMC thresholds. There is some evidence for Aβ40 and α-synuclein that higher monomer concentrations increase the order of the critical nucleus. This follows theoretically from (25), which suggest that a denser peptide environment in higher-order oligomers can have a detectable influence on β-propensity of their monomers [6]. It is supported by Monte Carlo simulations of oligomerization, where peptides are modeled as rigid rods [6]. Such a method predicts weak dependence of critical nuclei size inside oligomers which average size increases with concentration for weak β-prone peptides and comparable with oligomer orders for more β-prone [6]. However, concentration of suggested slope change for α-synuclein is closer to concentration of disaggregation of micelle-like clusters into oligomers rather to CMC [27], which agrees with (29) in case of decreased parasitic cluster size *N*_*off*_.

According to some thermodynamic models, longer peptides exhibit more favour towards oligomerized states [34]. However, in series of homopeptides, longer may demonstrate decrease in nucleation rate [35]. The formation of higher-order oligomers is favored by dominant interactions between aggregation-prone regions, outweighing the entropic cost of reduced degrees of freedom [34]. Kinetically, larger primary nuclei sizes in longer peptides may arise from their increased likelihood of anchoring and mutual unfolding. However, this rule is applicable only for peptides with similar sequences, and can not be applied even qualitatively for peptides with distinct secondary structures and charge distributions. The rate of primary nucleus formation is believed to be mainly governed by the peptide’s β-sheet propensity [6]. For instance, the calculations show that this property is not related to size of a nucleus. For instance, predicted β-propensities of insulin and *α*-synuclein are comparable, despite the first is more than two-fold shorter and contains nearly five times fewer helical residues as the second one. This suggests that both peptides possess short, similarly sized regions that initiate rearrangements into β-strands. However, among peptides with change in slope (Table 1), longer and more helical seem to be more prone to assemble into larger productive oligomers. PrP and α-synuclein have the longest helical regions, and the largest theoretical nuclei (the slope above CMC for α-synuclein). It can be explained by the fact that the long enough helical regions are stabilized by interchange between α- and π-helices [36], as has been approved computationally for α-synuclein [31] with stable helical up to tetramers. We propose that PrP and α-synuclein, due to their relatively extended sequences, form more stable hydrogen bond networks between adjacent monomers in fibrils, thereby reducing the likelihood of fragmentation and making the elongation (with capping) model more applicable. Insulin, α-synuclein, glucagon, and PrP posses significant helical fractions, about or more than 50%, and the strongest slope changes (except PrP with just the steepest slope among discovered peptides, and α-synuclein with too abrupt change not provided in the table), which also shows that helical peptides tend to assemble into higher-order oligomers [4], relatively small *N* ^*^ of glucagon for high concentrations (smaller than for lower) can be explained by 29, for example, *N*_*off*_ *≈* 2 transforms *N* ^*^ *≈* 2.16 (Table 1) into *N* ^*^ *≈* 4, resulting probably from mix of 2-, 3-, and 6-mers. For Sup35, only fragmentation model produced plausible *N* ^*^ *≈* 2, which is consistent with its known enhanced fragmentation in comparison with other peptides, including α-synuclein, slope of which corresponds to models without fragmentation [37].

Wider abundance of higher molecular mass oligomers does not necesseraly provide larger fibril nucleus size (Table 1), supporting off-pathway theory that many of lag-time oligomers are actually off-pathway peptide accumulators [1, 7]. However, it does not reject possibility of oligomerized monomers to be more prone to β-sheet conversion. Low-molecular (monomers, dimers, trimers) nuclei are readier described as supramolecular complexes (25) rather than forming new phases (24). Assumption of oligomer-wise assembly, when smaller oligomers merge into higher-order instead of monomer-wise attachment, is not supported by [7], and by our investigation with lower estimations of nuclei orders.

The biotechnological significance lies in the observation that hydrogen bond patterns, derived from known fibril structures and associated with higher aggregation propensity, do not necessarily correlate with faster aggregation kinetics. Stronger hydrogen bond networks may hinder the fragmentation of early amyloid strands, a factor omitted in prior studies predicting aggregation rates from peptide structures [40].

## 4 Conclusions

Double logarithmic dependence of lag times on peptide concentrations exhibits quick universal hands-on tool to estimate roles of oligomers of different sizes and secondary amyloid propagation pathways, and may help to unveil general properties like critical concentrations or nucleus sizes for whole groups of peptides. We hope it will advance our understanding of productive and inhibiting oligomers, and help to develop techniques for kinetic and thermodynamic escape of toxic species. Predicting the predominant nucleus sizes for a peptide based on its amino acid sequence remains challenging, as integral properties such as length and fraction of helical secondary structures are insufficient to fully determine the outcome. Future aims include improvement the model’s capacity to accommodate various aggregation pathways and predict theoretical structures of nuclei from the sequence.

Our calculations demonstrate that a reduced slope in the double logarithmic plot of lag time versus concentration may arise not only from smaller primary nuclei but also from enhanced fragmentation rates and fibril end-capping constants. A significant implication is that variations in these slopes among peptides may be primarily governed by secondary events rather than the size of the primary nucleus. The observed concavity of the lines can be explained not solely by the formation of higher-order oligomeric species, but also by the emergence of complex, branched nucleation-propagation pathways.

The examples show how two readily measurable experimental parameters, lag time and total peptide concentration, can be compared with thermodynamic parameters, such as equilibrium constants of association, derived from concentration-dependent spectral measurements, and computational methods like Monte Carlo simulations and molecular dynamics. Together, these insights should provide a comprehensive understanding of the association and aggregation mechanisms of a peptide of interest.

## Acknowledgements

This work was supported by the grant number 19-74-30007 of the Russian Science Foundation.

## Declarations

### Code availability

The complete program code is publicly accessible at https://github.com/ivankovlab/concentration_and_lag_time.

